# Early mechanisms of whisker development: Prdm1 and its regulation in whisker development and evolutionary loss

**DOI:** 10.1101/2021.03.11.433122

**Authors:** Pierluigi Giuseppe Manti, Fabrice Darbellay, Marion Leleu, Bernard Moret, Julien Cuennet, Frederic Droux, Magali Stoudmann, Gian-Filippo Mancini, Agnès Hautier, Yann Barrandon

## Abstract

Whiskers (vibrissae) are miniaturized organs that are designed for tactile sensing. Extremely conserved among mammals, they underwent a reduction in primates and disappeared in the human lineage. Furthermore, whiskers are highly innervated and their mechanoceptors signal to the primary somatosensory cortex, where a column of neurons called “barrel” represents each of them. This structure, known as barrel cortex, occupies a large portion of the somatosensory cortex of the rodent brain. Strikingly, *Prdm1* conditional knockout mice are one of the rare transgenic strains that do not develop whisker hair follicles while still displaying a pelage (Robertson et al. 2007). Here we show that *Prdm1* is expressed early on during whisker development, more precisely in clusters of mesenchymal cells before placode formation. Its conditional knockout leads to the loss of expression of *Bmp2, Shh, Bmp4, Krt17, Edar, Gli1* though leaving the β-catenin driven first dermal signal intact. Furthermore, we prove that *Prdm1* expressing cells not only act as a signaling center but also as a multipotent progenitor population contributing to the formation of the dermal papilla, dermal sheath and pericytes of the vascular sinuses of vibrissae. We confirm by genetic ablation experiments that the absence of motile vibrissae (macro vibrissae) formation reverberates on the organization of nerve wiring in the mystacial pads and organization of the barrel cortex. We prove that *Lef1* acts upstream of *Prdm1* and identify a potential enhancer (named Leaf) that might be involved in the evolutionary process that led to the progressive reduction of snout size and vibrissae in primates.

## Introduction

Whiskers (or vibrissae – from the latin *vibrio*) are exquisite miniaturized sensory organs specialized in tactile sensing (Petersen 2007), (Brecht 2007), (Diamond et al. 2008). Conserved throughout evolution, whiskers underwent a reduction throughout the primate adaptive radiation (Van Horn 1970) and disappeared completely in the human lineage; yet vestiges of whisker capsular skeletal muscles remain in the human upper lip (Tamatsu et al. 2007). Those specialized hair follicles are bigger both in length and width compared to pelage ones and are enveloped by vascular sinuses conferring rigidity to the hair shaft. Fibers of striated muscle have an insertion on the capsula and encompass the vascular sinuses. While macro vibrissae are motile and used for distance detecting/object locating, micro vibrissae are immotile and used for object identification.

The processing of whisker-acquired information occurs in the barrel cortex, where each whisker is represented by a discrete and well-defined cytoarchitectonic structure that goes under the name of barrel (Woolsey and Van der Loos 1970). The barrel map occupies a large area of the brain, it is in large part genetically specified and forms early on during development (Erzurumlu and Gaspar 2012). As whisker pattern is established earlier and independently from innervation, the hypothesis being that whiskers impose their own pattern onto the somatosensory cortex in the homeomorphic fashion has arisen (Andrés and Van der Loos 1985).

*Prdm1* is a zinc-finger transcriptional repressor (Keller and Maniatis 1991) that has been proven to be a master regulator controlling terminal differentiation of B-lymphocytes (Angelin-Duclos et al. 2000) (Shaffer et al. 2002) (Shapiro-Shelef et al. 2003); it also governs T-cell homeostasis (Martins et al. 2006) and primordial germ cells specification (Vincent et al. 2005), stem cell maintenance in the sebaceous gland (Ohinata et al. 2005) and skin differentiation (Horsley et al. 2006). *Prdm1* has also been shown to play a crucial role during whisker development (Robertson et al. 2007).

*Prdm1* (also known as *Blimp1*) conditional knockout mice are one of the very rare transgenic animals entirely lacking whisker (vibrissae) follicles while pelage hair follicles develop physiologically. The loss of this gene impairs whisker development (Magnúsdóttir et al. 2007), though the exact stage at which the development is halted has not been identified yet. Furthermore, it is still not known which type of mystacial vibrissae (macro- and/or micro-) are impacted by *Prdm1* knockout.

Moreover, Robertson et al. proved that *Prdm1* positive mesenchymal cells give rise to the mature dermal papilla and expand to form a mesenchymal layer immediately surrounding the hair follicles. However, what this mesenchymal layer gives rise to the adult whisker is a question that has not been answered yet.

Intriguingly, the reverberation of whisker development halt onto the barrel cortex has not been investigated yet. The re-organization of the somatosensory cortex is a phenomenon of great importance for evolutionary reasons, given the expansion of the brain areas dedicated to the processing of the sensory organs that evolutionary took over.

Eventually, little is known on the regulation of *Prdm1* during whisker development. Investigating this subject is of fundamental importance as the loss of potential regulatory elements in upstream genes might explain the reduction that occurred throughout the primate adaptive radiation and that led to reduction of snout size and vibrissae while hands and eyes of diurnal monkeys took over as sensory organs.

## Results

### Prdm1 is a master gene of whisker follicle development

We localized *Prdm1* expression by immunofluorescence in *Prdm1* mEGFP embryonic whisker pads during the first stages of whisker development (Figure 1A); those mice dispose of an mEGFP reporter recombined in frame with the exon 3 of *Prdm1* (Ohinata et al. 2005) (SFig 1A, Figure 1B). mEGFP expression in heterozygous embryos can be detected in a specific cluster of mesenchymal cells underlying the monolayer of embryonic epidermis (referred as stage 0 of whisker development, Hardy 1992) that will later on form the whisker epidermal placode. It continues to be expressed in this compartment until stage 4, when its expression is turned on in the inner root sheath (IRS) of the follicle. At stage 5, *Prdm1* expression disappears in the dermal papilla (Figure 1C). Those results are validated by *Prdm1* immunohistochemistry (IHC) in wild type whisker pads (Figure 1D).

**Figure 1.**
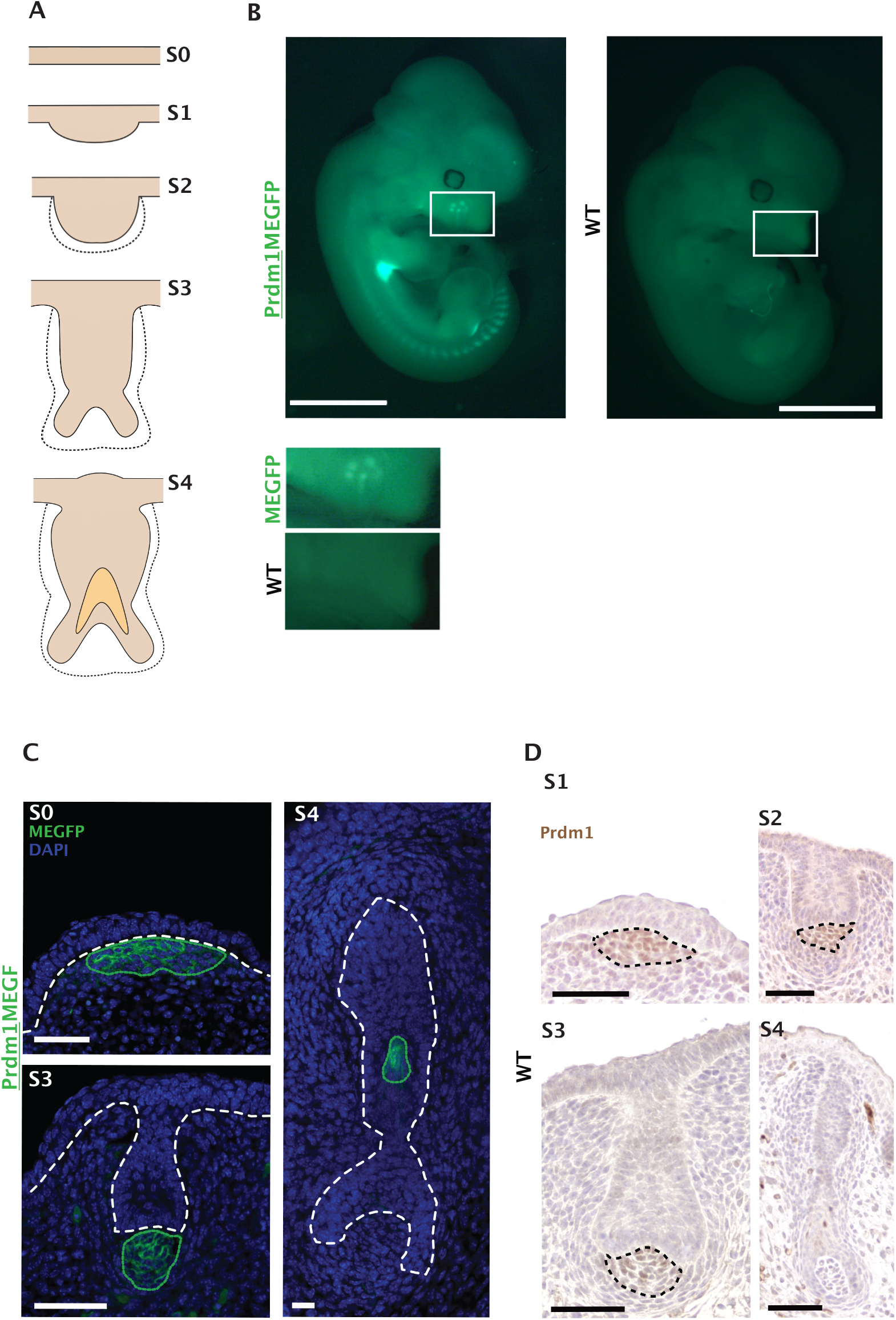
(A) Schematic representation of the whisker follicle developmental stages (S1-S4). Whisker development starts around embryonic day 12.5 (day 0 of whisker development) when an early dermal condensate appears in the whisker pad below the embryonic epidermis, which will in turn thicken to form a placode in stage 1 (Figure 5). Subsequently, an epidermal down growth (stage 2) and a dermal papilla (stage 3) are formed. A hollow cone (stage 4) develops by hardening of cells belonging to the hair matrix, thus giving rise to the inner root sheath. Image elaborated on (Hardy 1992). (B) *Prdm1*MEGFP versus wild type (wt) embryo at e12.5. Fluorescent signal is observed in the developing whisker pad, forelimb, hindlimb and somites. Right below full embryo pictures, magnification of the transgenic and wt whisker pad. (C) Prdm1 IHC on developing whisker pad. Left panel, *Prdm1* is expressed in the mesenchymal compartment from stage 1 (S1) to stage 3 (S3) of whisker development. Black dashed circles envelop the areas where *Prdm1* is expressed. (D) GFP immunofluorescence on *Prdm1*MEGFP whisker pads. On the top left, *Prdm1* can be detected before placode formation (S0). The GFP expression in the dermal fibroblasts at stage 3 (S3) and in the IRS at stage 4 (S4) confirms that the reporter mouse recapitulates the endogenous pattern of expression of *Prdm1*. The white dashed lines indicate the epidermal-dermal junction in S0-S3 and demarcate the follicle from the surrounding mesenchyme in S4. The green dotted lines indicate the areas of GFP expression in *Prdm1*MEGFP mice. Scale bar: 50 μm.

On the other hand, *Prdm1* expression in the dermal condensate of both head and back pelage hair follicles starts at embryonic day 14.5 (stage 0 of pelage follicle development) but is transient as it disappears at stage 2 (SFig 2A).

**Figure 2.**
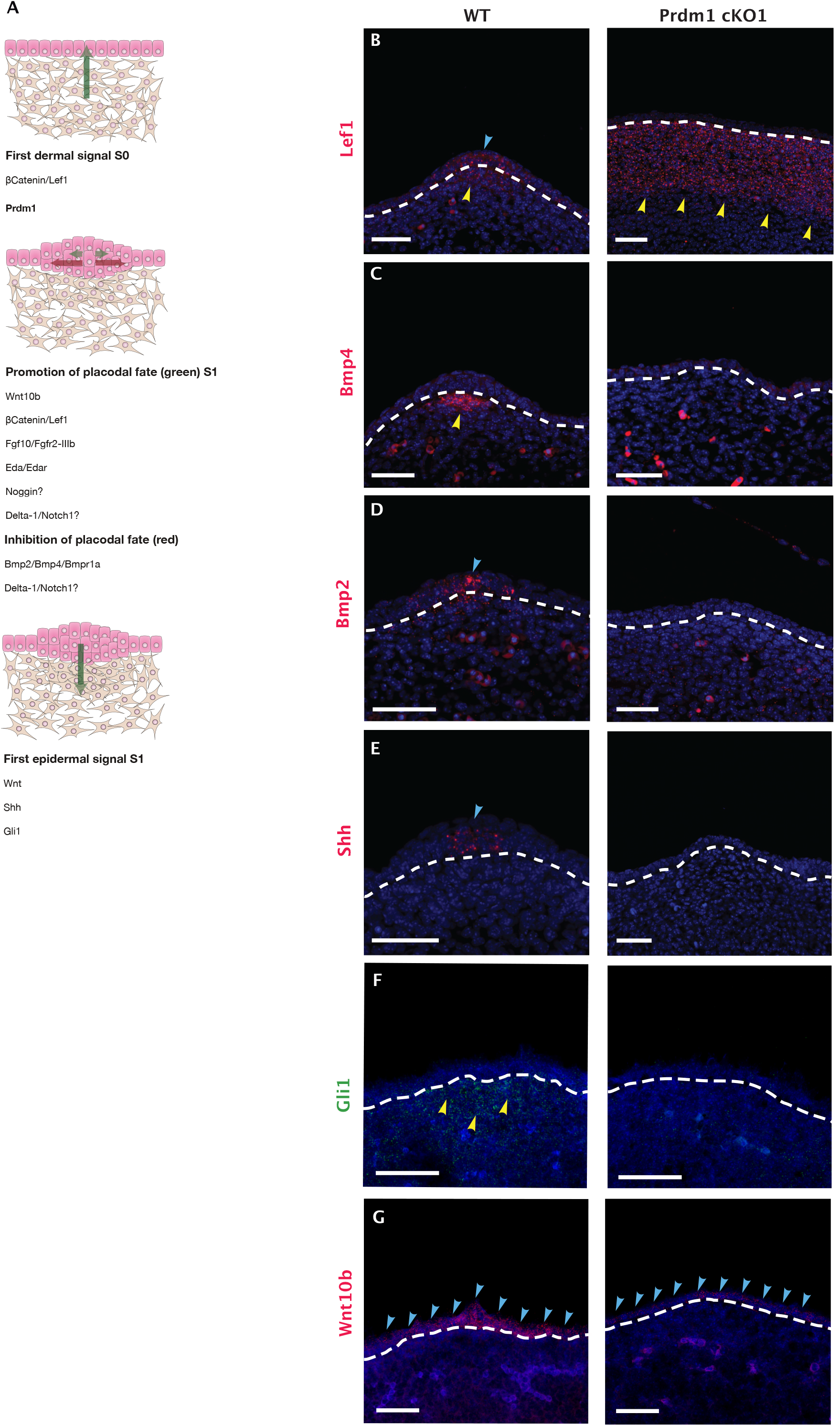
(A) Expression of early whisker developmental genes in the *Prdm1* cKO1 mouse. Molecular mechanisms underlying the early steps of whisker development. The dermis delivers a βcatenin based homogenous first signal to the overlying epidermis in order to initiate placode formation. The placode in turns both sustains their growth and inhibit the formation of other placodes in the surrounding epidermis. The promotion of the placodal fate is sustained by several molecules including Wnt10b, βcatenin/*Lef1*, Fgf10/Fgfr2-IIIb, Eda/Edar, Noggin, Delta-1/Notch1, whereas the inhibition is based upon *Bmp2*/Bmp4/Bmpr1a and Delta-1/Notch1. Thus, the placode conveys a first epithelial signal, leading to the clustering of the mesenchymal cells underneath into the dermal condensate. This process mainly relays on *Wnt* signaling and *Shh*. Image modified from (Millar et al. 2002). Fluorescent ISH on e12.5 wild type embryos (S1 of whisker development) reveals that, in physiological whisker development, *Lef1* expression is initially homogeneous in the mesenchyme and later confined to the epithelial placode and underlying mesenchyme (B); *Bmp2* marks the epithelial placode (C); *Bmp4* the underlying mesenchyme (D); *Shh* is expressed by the placode and induces the condensation of the mesenchyme (E); W*nt10b* expression is restricted in placodes (F); *Gli1* is upregulated in the pre-follicular mesenchyme. In e 12.5 cKO1 embryos, *Lef1* expression remains homogenous throughout the mesenchyme; *Bmp2, Bmp4, Shh* and *Gli1* are no longer detectable wile Wnt10b fails to be upregulated in placode areas; whisker follicles cannot thus reach stage 1 of whisker development. The dashed lines indicate the dermo-epidermal junction. Hybridization is marked with arrows (green arrows indicates expression in the placode, yellow ones in the mesenchyme). Scale bar: 50 μm.

To position *Prdm1* in the molecular cascade leading to whisker formation, we generated *Prdm1* knockout embryos. It has been previously shown that the constitutional knockout of *Prdm1* leads to severe impairment of the placenta resulting in early embryonic lethality. Therefore, we used a Sox2Cre deleter strain to bypass placental developmental halt (Vincent et al. 2005, see SFig1B). This transgenic line induces recombination in all epiblast cells by embryonic day (E) 6.5 but little or no activity in other extraembryonic cell types at this time (Hayashi et al. 2002). We were thus able to generate and harvest *Prdm1* homozygous conditional knockout embryos (cKO1) until E17, where exons four to eight are deleted by site-specific Cre-mediated recombination (SFig 1C).

We started by analyzing E12.5 and E13 cKO1 embryos, time-points when whisker follicle formation begins. Macroscopically, the cKO1 embryos are smaller compared to their wild type matches; as expected, they lack two or three digits (incomplete adactylia) in the forelimbs (SFig 2B). Microscopically, no whisker placode can be detected, and the dermal condensate formation cannot be observed (SFig 4). On the other hand, the development of pelage hair follicles is not impaired.

**Figure 3.**
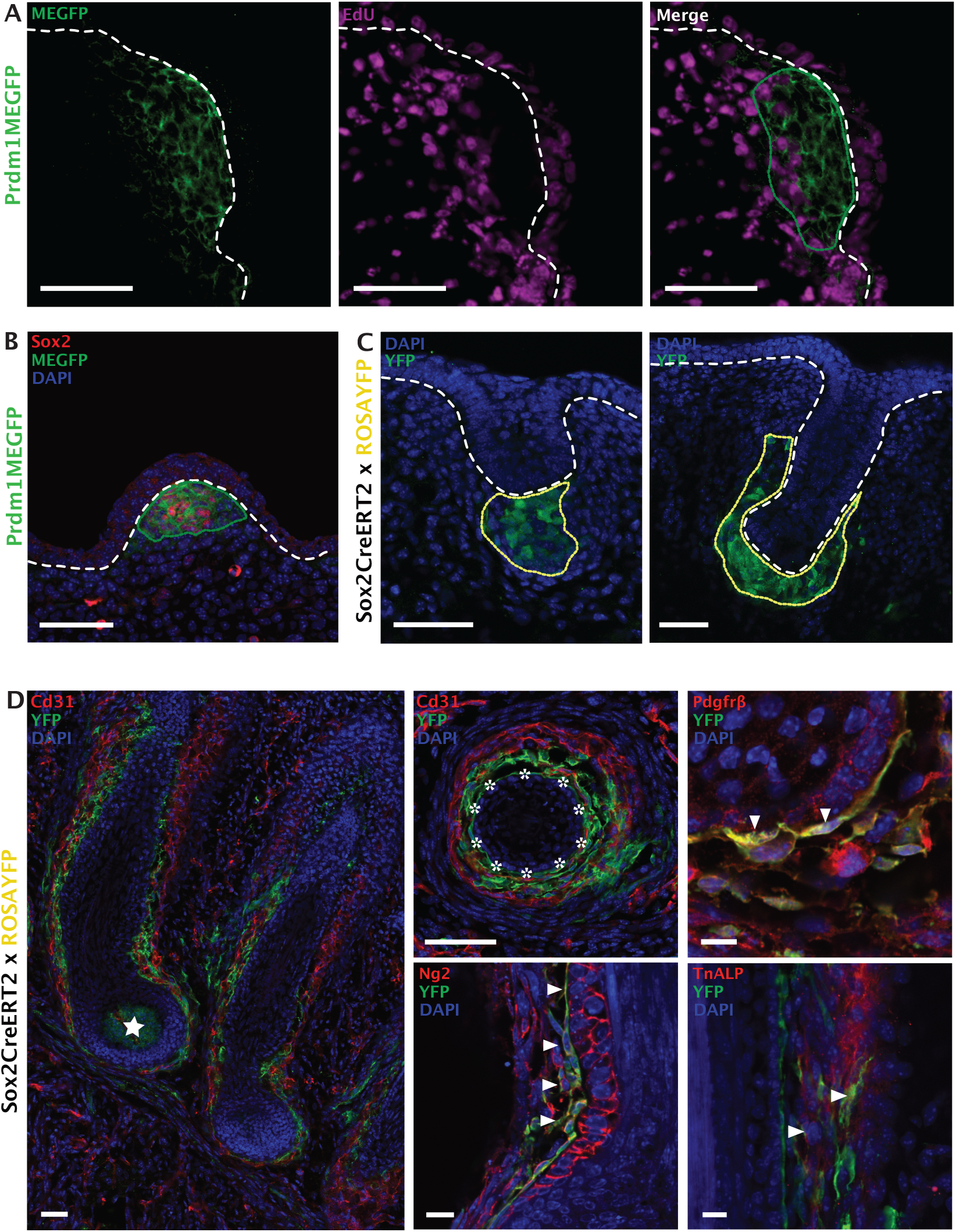
(A) The vast majority of GFP positive cells do not incorporate Edu contrarily to the peripheral ones at E12.5. (B) The Sox2 immunofluorescence on *Prdm1*MEGFP whisker pads reveals that *Sox2* marks a subpopulation of *Prdm1* positive cells. Note that Sox2 is also expressed in the putative oligodendrocytes surrounding the nerve endings surrounding the whisker pre-mesenchymal condensate. Cre expression was induced upon tamoxifen injection in E12.5 *Sox2*CreERT2/ROSAYFP embryos and examined for YFP expression either in the early stages (E13, E14) or at completion of development (E17, P3). (C) Note that *YFP* is first expressed in a cluster of mesenchymal cells right underneath the hair germ; when the latter becomes the hair peg, the YFP cluster envelops it into a mesenchymal cup. (D) Analysis at later time-points (E17-P3) reveals the extensive contribution of YFP+ cells to several lineages of the whisker follicle including the DP (starred) and the DS (asterisks). Several YFP+ cells in close contact with the endothelial ones (expressing Cd31) can be observed inside the vascular sinuses; those cells express markers of pericytes such as Tnap, Ng2, Pdgfrβ (arrows indicate areas of co-expression). White dashed lines (A, B, C) indicate epidermal-dermal junction. Green dotted lines indicate clusters of GFP positive cells (A, B); yellow dotted lines indicate the progeny of *Sox2* positive cells (YFP positive, C). Scale bars: 50 μm, 10 μm (Ng2), 20 μm (Tnap).

**Figure 4.**
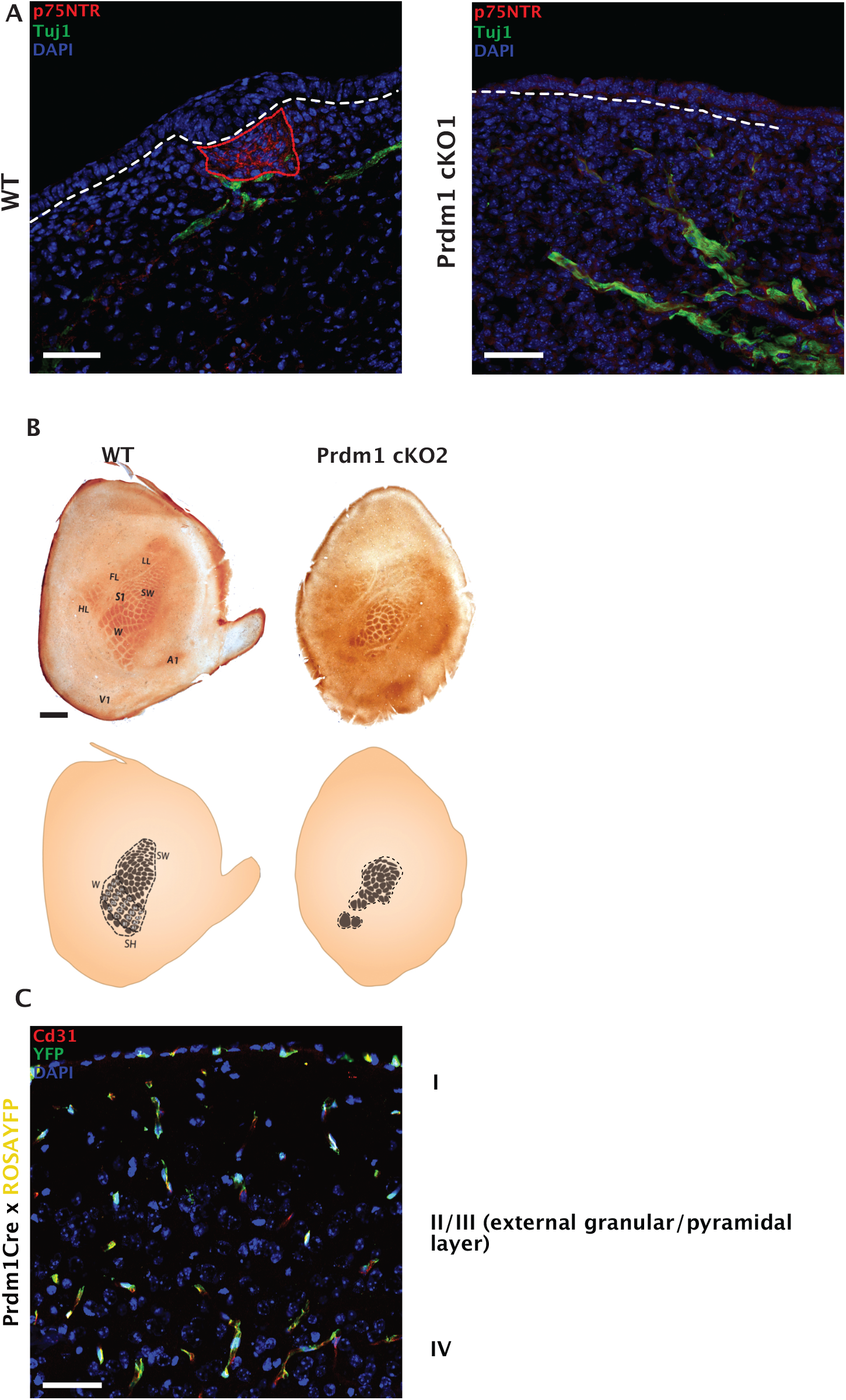
(A) Tuj1 immunofluorescence on *Prdm1* cKO whisker pad at E13.5 reveals the innervation process in the early stages of whisker development. Note that the nerve fibers encapsulate the mesenchymal condensate in the wild type whereas they act as free nerve endings in the cKO. (B) Cytochrome oxidase staining on the barrel cortex of *Wnt1*Cre driven *Prdm1* KO mice. Note the absence of the vast majority of barrels representing the macro vibrissae and the rearrangement of the ones representing the micro vibrissae. (C) Cd31 (red) and GFP immunofluorescence on the somatosensory cortex of *Prdm1*Cre/ROSAYFP mice. Note that the axons and neurons of layer four of the cortex have never expressed *Prdm1* and that YFP+ positive cells are of endothelial origin. The red dotted line in A indicates the area of dermal condensate expressing p75NTR. Scale bar: 50 μm.

To understand where *Prdm1* stands in the molecular cascade leading to whisker formation (Figure 2A) we looked with fluorescent *in situ* hybridization (FISH) at the expression of the genes involved in the first molecular steps of hair follicle morphogenesis both in the epithelial and mesenchymal compartment. β-catenin is implied in establishing the first dermal signal (Noramly et al. 1999); consistent with this, *Lef1* is expressed in the mesenchyme of the mouse vibrissa pad prior to vibrissa follicle development, and initiation of vibrissa follicle development is dependent on its expression (van Genderen et al. 1994), (Kratochwil et al. 1996). In E13 homozygous cKO1 mice, *Lef1* is expressed homogeneously in the mesenchyme of the whisker pad, thus indicating that the first signal is intact; however, the physiological restriction of expression of *Lef1* in the whisker placode and underlying mesenchyme does not occur (Figure 2B).

The gene cascade that is activated in placode formation is not activated in *Prdm1* cKO1 mice. No placode formation can be observed both macroscopically and microscopically. The patterned upregulation of molecules involved in the promotion and inhibition of placodal fate including *Bmp4* (Figure 2C) in the pre-follicle mesenchyme and *Bmp2* (Figure 2D) in the epithelial compartment does not occur. The same applies for a molecule involved in the first epidermal signal, SHH – secreted by the placodal cells (Figure 2E). *Gli1* is not upregulated in the pre-follicular mesenchyme (Figure 2F). We found that *Wnt10b* is diffusely expressed in the monolayer of epidermal cells in the sites of whisker formation in cKO1 and does not show a marked upregulation in placodes (as in wild type whisker pads) (Figure 2G). Furthermore, RT-qPCR on whisker pads shows a statistically significant decrease of *Edar*, involved in the promotion of placode, and *Keratin 17*, whose expression arises within the single-layered, undifferentiated ectoderm of embryonic day 10.5 mouse fetuses giving rise, in the ensuing 48 hours, to the epidermal placodes (McGowan and Coulombe 1998) (SFig 4). Intriguingly, *Lef1* is expressed uniformly in the mesenchyme of cKO1 embryonic whisker pads, and the typical restriction to the whisker placode and pre-follicle mesenchyme does not occur.

### Whisker inducing mesenchyme contributes to the formation of both the dermal sheath and capsula of adult mystacial whisker follicles

To investigate the proliferative activity of *Prdm1* expressing cells, we administered a short pulse (2 hours) of nucleotide analog ethynyldeoxyuridine (EdU) just prior to analysis to *Prdm1*MEGFP pregnant mice carrying E12.5 embryos. We could observe that *Prdm1* expressing cells can be classified into two subpopulations: the quiescent ones - contiguous to the embryonic epithelium - and the proliferative ones at the periphery (Figure 3A).

We then aimed at identifying the progeny of *Prdm1* expressing cells in the whisker follicle by means of lineage tracing. By crossing the *Prdm1*Cre transgenic strain with ROSAYFP mice (SFig 1D, E), we were able to follow the progeny of *Prdm1* expressing cells at different time points during embryonic development (SFig 5).

**Figure 5.**
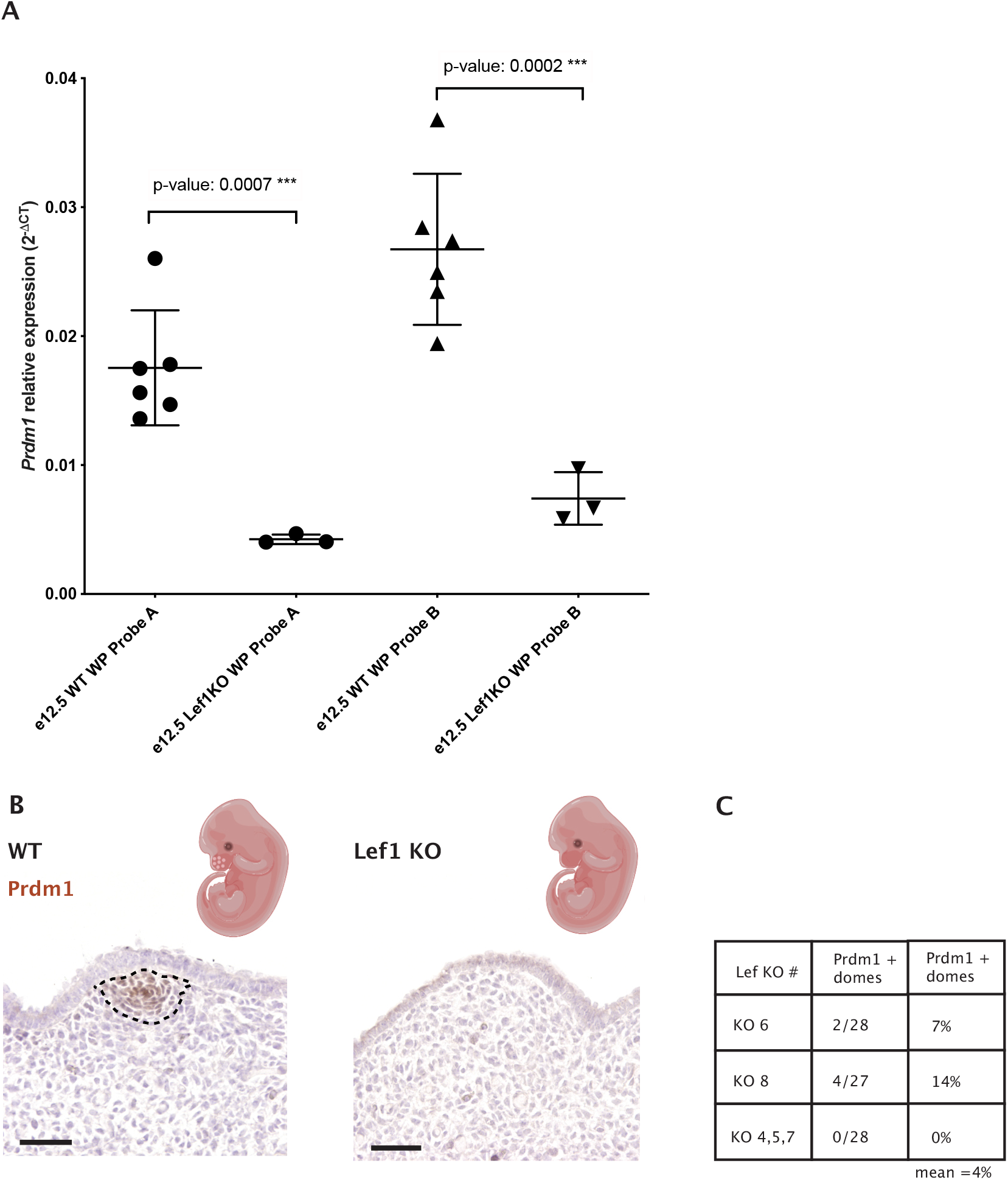
Quantification of *Prdm1* by RT-qPCR in both wild type and *Lef1* KO E12.5 whisker pads (each dot represents a sample) indicates a severe decrease of *Prdm1* expression in *Lef1* KO mice compared to the WT counterpart (p-value:0.0007***) (B) *Prdm1* IHC on WT and *Lef1* KO whisker pads (E12.5). Note the absence of expression of *Prdm1* in the ectodermal elevation preconfiguring sites of whisker induction in the *Lef1* KO embryos (KO 4, 5, 7). (C) Quantification of *Prdm1* expression in *Lef1* KO whisker pad (D) Schematic representation of the molecular mechanisms leading to whisker hair follicle formation. (i) A first dermal signal is needed to induce placode formation in the overlying epidermal monolayer (ii) The epithelial placode forms relaying on signals that promote its growth and inhibit placode fate in the surrounding epithelial cells (iii) The placode produces the first dermal signal.

At E12.5, YFP expressing cells are located in the whisker mesenchymal condensate recapitulating the GFP expression in the *Prdm1*MEGFP embryonic whisker pad. However, at E13.5 it can be observed that the population of YFP positive cells expands and encompasses the area of mesenchyme surrounding the whisker hair germ. *Prdm1* can be detected in this area (except for the precursors of the dermal papilla) in whisker pad sections of both *Prdm1*MEGFP and WT mice at E12.5 and E13.5 (Figure 3B).

Because of *Prdm1* expression in both the endothelium and somites, the lineage tracing experiment could not be performed at a time point later than E13.5. However, we observed that Sox2 is co-expressed with *Prdm1* at E12.5 in the whisker mesenchymal condensate (Figure 3B). Consequently, we crossed *Sox2*CreERT2 and ROSAYFP mice and injected tamoxifen in pregnant females at E12.5 (SFig 1E, F); double transgenic embryos were analyzed at E14.5, E17.5 and P1-3. 48 hours after tamoxifen injection, YFP positive cells were detected in the dermal condensate of the hair germs. As the epidermal down growth proceeds, the mesenchymal YFP cells progressively encapsulate it (Figure 3C). When the whisker follicle reaches its final anatomical configuration, YFP is expressed by the whole dermal papilla (DP), dermal sheath (DS) and abundantly by cells residing into the vascular sinuses. More precisely, the latter are enmeshed with CD31 positive cells and display a perithelial position: both the latter and the expression of markers such as *Ng2, Pdgfrβ, Tnap* (Tissue nonspecific alkaline phosphatase) indicate that they are pericytes (Figure 3D).

### *Prdm1* genetic ablation leads to the disorganization of the rodent barrel cortex

To study the impact of whisker loss on the nervous system, we have generated cKO1 embryos and looked at their innervation in the developing whisker pad. In the wild type whisker pad, the afferent branches of the infraorbital nerve (ION) encapsulate the whisker mesenchymal condensate without penetrating it; on the contrary, they terminate as free nerve endings in the KO counterpart (Figure 4A).

To evaluate the consequences of the impaired whisker innervation on the nervous system, we have adopted the *Wnt1*Cre as a deleter strain in order to bypass the neonatal lethality that occurs in cKO1. *Wnt1* is expressed virtually in all neural crest derivatives, including the whisker pad mesenchyme and, especially, the mesenchymal compartment surrounding the developing whisker follicle. *Wnt1*Cre driven *Prdm1* conditional KO mice (cKO2 – see SFig 1E) are viable and lack almost all the macro vibrissae except for the 1-3 distal ones of the first row, as observed both macroscopically and microscopically (SFig 6).

**Figure 6.**
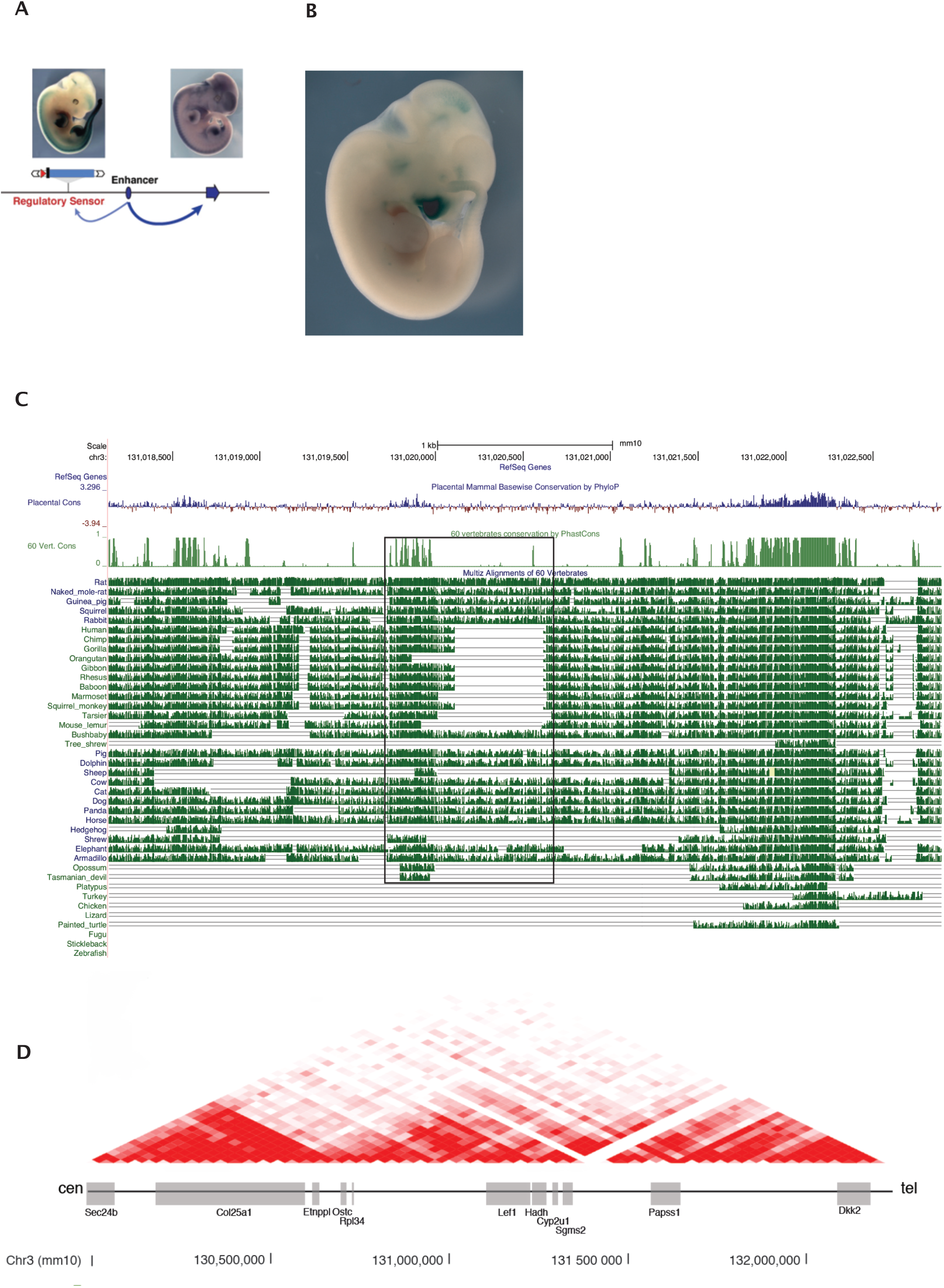
(A) The TRACER database pinpoints that the regulatory region of *Lef1* resides centromeric to his promoter and that is active in e11.5 whisker pads. (B) *Lef1* is active in the whole e11.5 whisker pad, coherently with the homogeneous expression of the first dermal signal. (C) Multispecies alignment showing a primate specific deletion of a sequence adjacent to a conserved region. Species analyzed include animals with developed whiskers (i.e. rat, cat, dog, rabbit, guinea pig, squirrel, horse, naked mole rat, pig), primates (human, chimp, gorilla, gibbon, rhesus, baboon, squirrel money, orangutan, marmoset, tarsier) and the human lineages. (D) HiC profile around *Lef1* in murine ES cells (Dixon et al. 2012). Cen: centromeric. Tel: telomeric

We retrieved the brains of both WT and cKO2 mice after p21 to account for the developmental maturation of the somatosensory system and sectioned their flattened somatosensory cortex tangentially to visualize the organization of the barrel cortex (Figure 4B). The cytochrome oxidase staining revealed that cKO2 barrel cortex undergoes a major rearrangement; the residual macro vibrissae are represented by enlarged barrels; the barrels corresponding to the micro vibrissae are, however, still present, even though their pattern is highly disorganized (Figure 4C).

To exclude the expression of *Prdm1* in the developing barrel cortex - and thus that the barrel cortex phenotype can be ascribable to the loss of *Prdm1* in the nervous system - we crossed *Prdm1*Cre with ROSAYFP mice and looked at YFP expression in the cerebral cortex (Figure 4D). YFP is expressed only by endothelial cells and not by the thalamocortical axons or by layer 4 cortical neurons, thus excluding the aforementioned possibility.

### *Lef1* acts upstream of *Prdm1* in the whisker hair follicle developmental cascade

We reasoned that β-catenin/Lef1 might act upstream of *Prdm1* during whisker follicle development. To prove this, we obtained *Lef1* constitutional KO embryos by crossing homozygous *Lef1*^tm1Rug^ mice (see SFig1G) (Van Genderen et al 2004) and investigated *Prdm1* expression in the E12.5 whisker pad both at the mRNA and protein level.

At a molecular level, the quantity of *Prdm1* transcript in the *Lef1* KO embryos is lower compared to the wild type counterpart, as demonstrated by RT-qPCR (Figure 5A). We sectioned the E12.5 whisker pads from *Lef1* KO and WT mice and localized *Prdm1* by IHC, focusing on the mesenchymal cells located under the characteristic surface elevations of the whisker pad that constitute the sites of whisker placode induction – from now on referred as areas of analysis. We analyzed the transverse sections of the embryonic whisker pad through (a) the primitive nasal cavity (b) the vomeronasal organ and (c) the tongue of five *Lef1* KO embryos. Out of five *Lef1* KOs, three areas did not express *Prdm1*. As for the remaining two embryos, we could observe *Prdm1* expression in 2/28 and 4/27 areas (Figure 5B and 5C).

To understand if the de-regulation of *Prdm1* and/or *Lef1* might help to explain vibrissae reduction and their eventual loss in humans, we looked at their regulatory regions. As for *Lef1*, a transposable enhancer trap mapping to a particular locus (chr3:130,927,182-130,927,529 in mm10 assembly) suggests that its regulatory region is located on the centromeric side of the gene, in the adjacent gene desert (TRACER LacZ expression database, SB line name 183038-emb20) (Chen et al. 2013). The transgene expression can be observed at E11.5 in the brain, mammary glands, whisker pad and the tip of tail, tissues/organs where *Lef1* is physiologically expressed during development (Figure 6A).

We compared the multispecies alignment of animals with and without whiskers in the aforementioned locus searching for putative regulatory elements. Conservation scored by PhastCons indicated the presence of an element (878bp) conserved throughout the two categories of species (chr3:131,019,746-131,020,624) which contains a sub-region (521bp) specifically absent in animals deprived of functional whiskers (human, chimp, gorilla, gibbon, rhesus, baboon and squirrel monkey) mapping chr3:131,020,103-131,020,624 (mm10 assembly) (Figure 6B). Altogether, this DNA element (Leaf, chr3: 131,019,746-131,020,624) is located in the same topological associating domain (TAD) where *Lef1* resides (Figure 6C).

To observe if this region can contact the promoter of *Lef1*, we performed 4C-Seq. The analysis was conducted on whole microdissected whisker pads at E12.5 using primers positioned in the promoter of *Lef1*. Adult kidney was used as the negative control as *Lef1* is not expressed in this tissue (Figure 7A and 7B, SFig 7, Darbellay and Necsulea, 2020).

**Figure 7.**
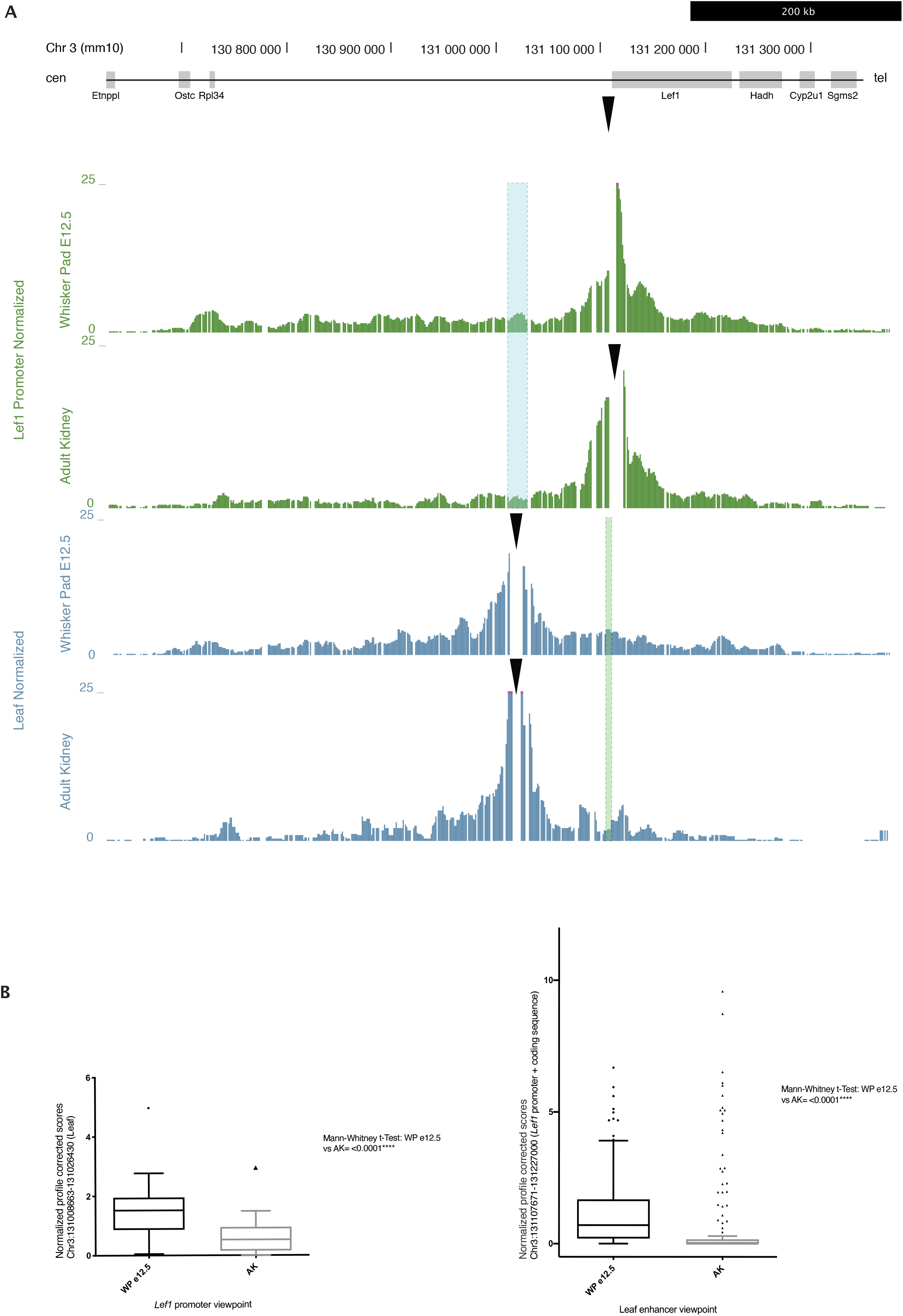
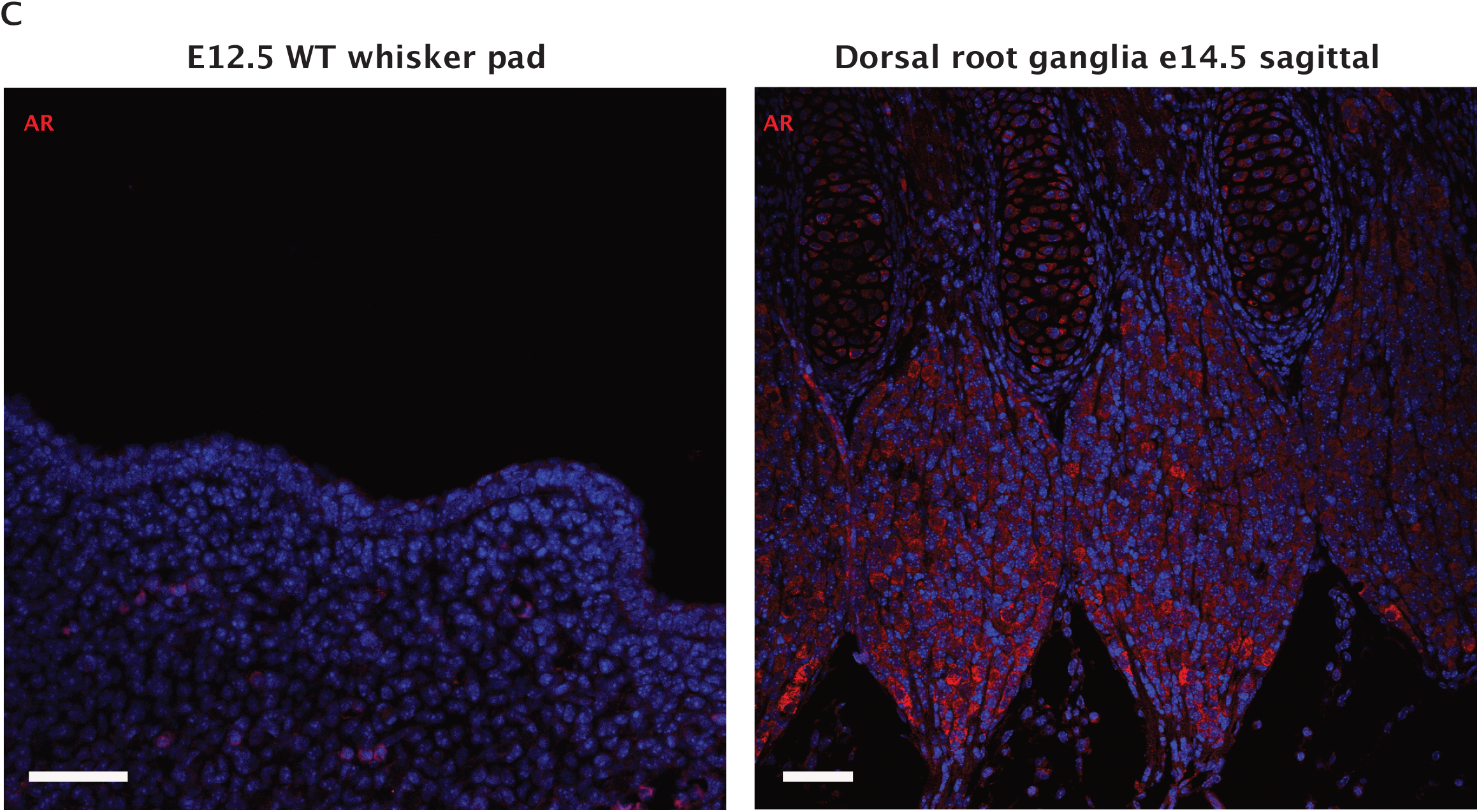
(A) 4C-Seq on e12.5 whisker pad and adult kidney (negative control). In green, the *Lef1* promoter (viewpoint, black arrow) contacts Leaf more frequently in the e12.5 whisker pad compared to the adult kidney. In blue, Leaf (viewpoint, black arrow) has a higher number of contacts on *Lef1* promoter and coding sequence in e12.5 embryos compared to the adult kidney. Tracks are normalized (for profile-corrected tracks see Supplementary Figure 6). (B) Normalized, profile corrected scores of *Lef1* promoter viewpoint on Leaf and vice versa show a statistically significant contact between the e12.5 whisker pad and adult kidney. (C) AR IHC shows no expression in the E12.5 whisker pad compared to dorsal root ganglia (positive control).

We found that *Lef1* promoter scores contact mainly in cis, within an area of around 700 kb surrounding the *Lef1* locus, corresponding to its TAD. More specifically, the centromeric region contains most of the peaks of interaction, whereas telomeric contacts occur chiefly with the coding sequence of *Lef1*, extending until the end of the coding sequence of the neighboring gene Hadh.

In the E12.5 whisker pad, the promoter of *Lef1* contacts several centromeric regions, among which the one containing the primate-specific deletion (Leaf). We proceeded to quantify the number of contacts that the *Lef1* promoter establishes with Leaf in the E12.5 whisker pad and the adult kidney both in the normalized and profile corrected dataset; we represented the sum of the fragments through boxplots. We found high statistical significance between the scores mapping Leaf in the E12.5 whisker pad compared to the adult kidney (****, Mann-Whitney test, p=0.0008 in normalized and p<0.0001 in PC).

In the reciprocal experiment, we looked at the contacts from the murine Leaf enhancer with the promoter and coding sequence of *Lef1*. We found high statistical significance between the scores mapping *Lef1* in the E12.5 whisker pad compared to the adult kidney, suggesting productive long-range interactions between Leaf and *Lef1* in the whisker pad (****, Mann-Whitney test, p<0.0001 in normalized and p<0.0001 in PC), thus confirming what observed in the specular viewpoint.

It has been recently demonstrated that a specific enhancer of the androgen receptor (Ar) is expressed in the whisker mesenchyme during development (McLean, 2011); the authors propose that the loss of this enhancer is associated to the loss of both sensory vibrissae and penile spines in the human lineage. Notably, the castration or genetic deletion of the Ar results in a reduced growth of whisker follicles in mice without their full disappearance.

To understand if the AR is involved in the early steps of whisker development together with *Prdm1* and *Lef1*, we performed both IHC and RT-qPCR on embryonic whisker pads at E12.5. The AR IHC shows no signal either in the whisker placodes or in the mesenchyme underneath it at E12.5; the RT-qPCR confirmed the absence of expression in the micro-dissected whisker pads at E12.5 and E13.5 (Figure 7C), thus excluding the hypothesis of its early involvement in whisker formation.

## Discussion

Overall, our results prove that (i) *Prdm1* acts at the level of the first dermal signal during whisker development and its expression is regulated by β-catenin/*Lef1* (ii) *Prdm1* is fundamental for the proper functioning of the whisker signaling center that will later contribute to the formation of the adult structures of the whiskers, more specifically the dermal sheath, the dermal papilla and the pericytes (iii) the disappearance of macro vibrissae induced by the genetic ablation of *Prdm1* causes the reorganization of the entire murine barrel cortex (iv) *Lef1* is positioned upstream of *Prdm1* in the whisker inductive cascade and the loss a putative regulatory element (Leaf) might contribute to the multi-step process that lead to the progressive de-functionalization and loss of whisker in primates and humans.

*Prdm1* has already been described as a key gene for whisker development (Robertson et al. 2007). We prove that *Prdm1* is expressed in clusters of mesenchymal cells before whisker placode appearance, thus implying a very early role in whisker formation. Our in-situ hybridizations on the *Sox2*Cre driven *Prdm1* KO confirm and expand the previous results of Robertson et al. We prove that in *Prdm1* cKO1 whisker pads the β-catenin based first dermal signal is intact; however, no placode formation occurs – as shown by *Wnt10b, Gli1, Edar, Krt18* ISH - and the first epidermal signal cannot occur. Thus, we can conclude that *Prdm1* operates at the level of the first dermal signal.

The fate mapping study conducted by Robertson et al. (Robertson et al. 2007) indicates that *Prdm1* expressing cells contribute to the formation of the DP and that the cells failing to be incorporated in the latter migrate to surround the hair shaft. To understand the fate of those specific cells, we performed a lineage tracing study using the *Sox2*CreERT2 strain, once proven the co-expression of *Prdm1* and *Sox2* in the cluster of mesenchymal cells right underneath the pre-placodal epithelium. Our results clearly illustrate that the population of cells expressing *Prdm1*/Sox2 gives rise to several lineages of the adult whisker, thus representing a population of multipotent progenitors. As expected, the mesenchymal pre-condensate contributes to the formation of the dermal papilla; interestingly we demonstrate that it also gives rise to the dermal sheath of the follicle, thus explaining the common inductive properties of the DP and DS (Oliver 1966), Eventually, we prove that it gives rise to pericytes residing in the whiskers’ vascular sinuses; their lineage is confirmed by their anatomical position and the expression of specific markers.

Robertson et al. shows that *Prdm1* expressing cells display no Ki67 expression and are thus not cycling. Our Edu uptake experiments show that the vast majority of them – the ones that are the most proximal to the placode – do not incorporate Edu, thus confirming Robertson’s data. However, we can observe that *Prdm1*/*Sox2* positive peripheral cells are proliferative. We speculate that peripheral cells progressively lose *Prdm1* expression and are thus able to re-enter cell cycle, whereas those underneath the placode retain its expression, functioning as signaling center during development and in the end constituting the dermal papilla of the mature follicle.

We further investigated the impact of *Prdm1* knockout on the murine nervous system. In cKO1 whisker pads, the afferent branches of the infraorbital nerve do not organize into plexuses surrounding the mesenchymal condensates but terminate as free nerve endings. Axonal guidance is most probably absent as for the lack of nerve guidance molecules secreted by *Prdm1* positive cells. The *Wnt1*Cre driven *Prdm1* KO (cKO2) mice lack almost all macro vibrissae though retaining micro vibrissae. This result suggests that *Prdm1* plays a key role in macro vibrissae development; on the contrary, the formation of micro vibrissae in cKO2 indicates that the signals required to induce their formation do not rely on *Prdm1* and still need to be investigated. Those conditional knockout mice have major rearrangements in the barrel cortex concerning also the representation of micro vibrissae with relevant evolutionary consequences.

Given that the first dermal signal is Wnt-based and that *Lef1* plays a crucial role in early whisker development, we decided to focus on the relationship between *Lef1* and *Prdm1*. The downregulation of *Prdm1* in *Lef1* KO whisker pads both at the mRNA and protein level clearly proves that the β-catenin-based uniform first dermal signal is indispensable to induce *Prdm1* expression. In our model, *Prdm1* is dependently arising on the first dermal signal insurgence and as stated above, is fundamental for placode formation. We reasoned that the loss of regulatory sequences in *Prdm1, Lef1* or both might explain the multistep evolutionary process that led in the first step to whisker de-functionalization and then disappearance. King and Wilson (Jahoda and Oliver 1984) postulated that “regulatory mutations account for the major biological differences between chimp and human”; it has already been elegantly demonstrated how enhancers regulate the craniofacial morphology (Attanasio et al. 2013). Intriguingly, it has been demonstrated how species-specific sequence changes in an evolutionary conserved enhancer of Shh contribute to its functional degeneration in snakes (Kvon et al. 2016), highlighting the prominent role of enhancers in morphological evolution.

It has been previously been shown that the spatial and temporal control of *Lef1* expression depends on different regions of the *Lef1* locus (Liu et al. 2004). A 2.5 kb segment of the human promoter can drive *LacZ* expression only in the mesenchymal compartment of the developing whisker follicles; furthermore, it contains a specific element responsive to *Wnt3a* and β-catenin (Liu et al. 2004). Furthermore, the congenital agenesis of molar number 3 (am3) in a murine strain is attributable to a locus mapping the 130.73-131.69 Mb region of chromosome 3, suggesting that *Lef1* is the strongest candidate for am3 (Shimizu et al. 2013).

We searched the TRACER database to find the regulatory region of *Lef1*; the transposon-based enhancer trap indicates that the latter resides centromeric to *Lef1* and that is active during whisker morphogenesis at E11.5. We identified a deletion in the regulatory region of *Lef1* that is specific to several primates (including human, chimp, gorilla, gibbon, rhesus and baboon) flanking a well-conserved region among many mammals. Leaf falls in the regulatory region identified with the TRACER database and in the TAD where *Lef1* resides. We thus postulated that this genomic sequence might have a putative enhancer activity in whisker development and, by modulating *Lef1* expression in the whisker pad, and indirectly *Prdm1*), might be involved in whisker disappearance during evolution. The 4C-seq analyses we performed show that Leaf is contacted by the *Lef1* promoter more often in the E12.5 whisker pad (embryonic age when whisker formation starts) compared to control tissue, thus suggesting a tissue-specific interaction. Those results are corroborated by the quantitative analysis of the scores mapping Leaf in the E12.5 whisker pad (and Lef1 in the reciprocal experiment) compared to the adult kidney, clearly underlining a highly statistically significant contact between Leaf and Lef1 promoter and vice-versa occurring specifically in the developing whisker pad. The function of this regulatory element is to be further investigated at the functional level.

When investigating AR expression in the whisker placodes at early stages, we could not detect its presence both at the RNA and at the protein level in both placode and underlying mesenchyme. We can thus rule out an early role of the enhancer that McLean and colleagues has described as involved in whisker loss in humans (McLean et al. 2011). Noteworthy is the fact that 2 strains of AR knockout mice generated by two independent groups (Yeh et al. 2002) (De Gendt et al. 2004) do not display whisker loss. Being it expressed in the mesenchymal compartment at later stages of hair follicle development, we hypothesize that it might play a role more in the de-functionalization of whiskers.

While great apes have lost macro vibrissae though retaining the micro vibrissae on lips (a phenotype that recalls our *Prdm1* cKO2 mice), cheeks and eyebrows, humans are the only known mammals to have lost both, though vestiges of vibrissal capsular muscles have been identified in the human upper lip (Tamatsu et al. 2007). In the model we envisage, whisker loss is a multistep process that started during the divergence of the species and that relies on the loss of tissue-specific regulatory elements of several genes involved in whisker formation; more specifically, the loss of Leaf might have contributed to the downregulation of *Lef1* expression in the snout of primates, thus contributing to whisker loss together with other mechanisms that have to be identified yet.

## Supporting information

SFig1

SFig2

SFig3

SFig4

SFig5

SFig6

## Figure Legends

**SupplementaryFigure 1**

Schematic representation of transgenic mouse lines used for this study.

**Supplementary Figure 2**

(A) Prdm1 IHC on developing head/back pelage hair follicles. *Prdm1* expression in pelage hair follicle development is transient, lasts until hair germ formation (S2) and is independent of the embryonic origin of the mesenchyme. Dashed circles envelop the areas where *Prdm1* is expressed. Scale bar: 50 μm. (B) WT e12.5 embryo compared to *Prdm1* cKO1. cKO1 embryos lack 2/3 digits and display no whisker placodes macroscopically.

**Supplementary Figure 3**

(A) Expression of Sox2 in wild type and *Prdm1* cKO1 whisker pad at e13.5. No dermal condensate can be detected in cKO1 embryos. Scale bar: 500 μm (B) Edar and Krt17 real time qPCR on E13 whisker pads of wild type and cKO1 mice.

**Supplementary Figure 4**

*Prdm1*Cre lineage tracing in developing whisker pads. *Prdm1*Cre was crossed with a ROSAYFP strain and developing whisker pads were analyzed at e12.5 and e13. YFP positive cells encompass the developing whisker. Scale bar: 50 μm.

**Supplementary Figure 5**

*Wnt1*Cre driven homozygous *Prdm1* knockout mice (cKO2) lack almost all the macro vibrissae except for the 1-3 distal ones of the first row, as observed both macroscopically and microscopically (C,D,E) compared to wild type animals (A,B). The barrel cortexes of the correspondent animals were stained with cytochrome oxidase and reveal a major rearrangement. Residual macro vibrissae are represented by enlarged barrels; the barrels representing micro vibrissae are still present, though highly disorganized.

**Supplementary Figure 6**

Profile-corrected tracks of 4C-Seq on e12.5 whisker pad and adult kidney (negative control). In green, the Lef1 promoter (viewpoint) contacts Leaf more frequently in the e12.5 whisker pad compared to the adult kidney. In blue, Leaf (viewpoint) has a higher number of contacts on *Lef1* promoter and coding sequence in e12.5 embryos compared to the adult kidney.

**Supplementary table 1.**
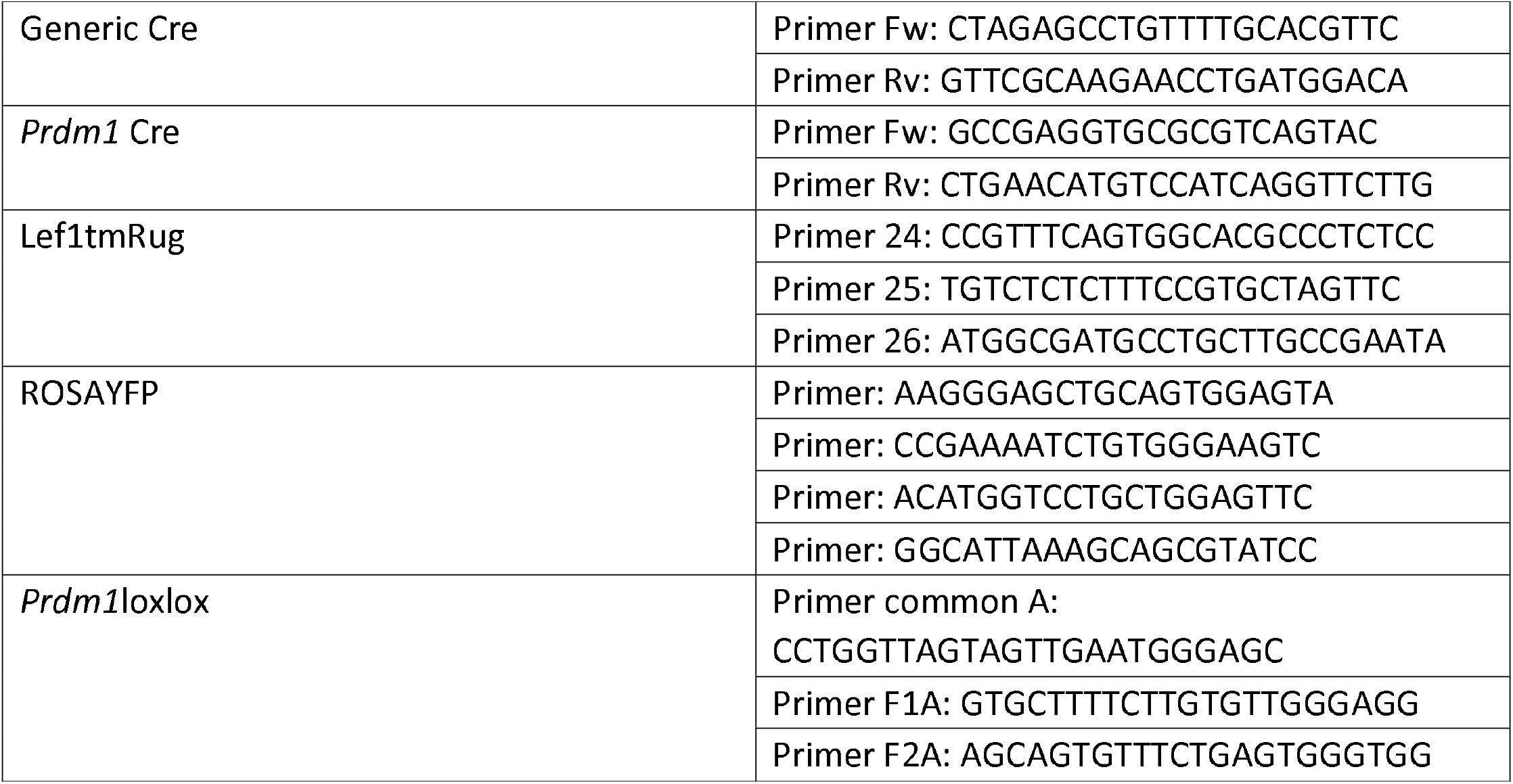
– List of genotyping primers

**Supplementary table 2.** – List of antibodies and dilutions

| Antibody | Species | Dilution | Clone | Company |
| --- | --- | --- | --- | --- |
| Prdm1 | Mouse | 1:500 | 3H2-E8 | Abcam |
| Sox2 | Mouse | 1:2000 | 9-9-3 | Abcam |
| p75 | Mouse | 1:1000 | 9G395 | UsBiologicals |
| GFP | Goat | 1:1000 | 6673 | Abcam |
| CD31 | Rat | 1:500 | MEC13.3 | BD Biopharmingen |
| Ng2 | Rabbit | 1:300 | AB5320 | Millipore |
| AR | Rabbit | 1:5000 | PG-21 | Millipore |
| Tuj1 | Goat | 1:300 | MMS-435P | Covance |
| Tuj1 | Rabbit | 1:300 | MRB-435P | Covance |
| Pdgrβ | Rabbit | 1:300 | 28E1 | Cell Signaling |
| GFP | Goat | 1:1000 | NB100-NB1770 | Novus |

**Supplementary table 3.**
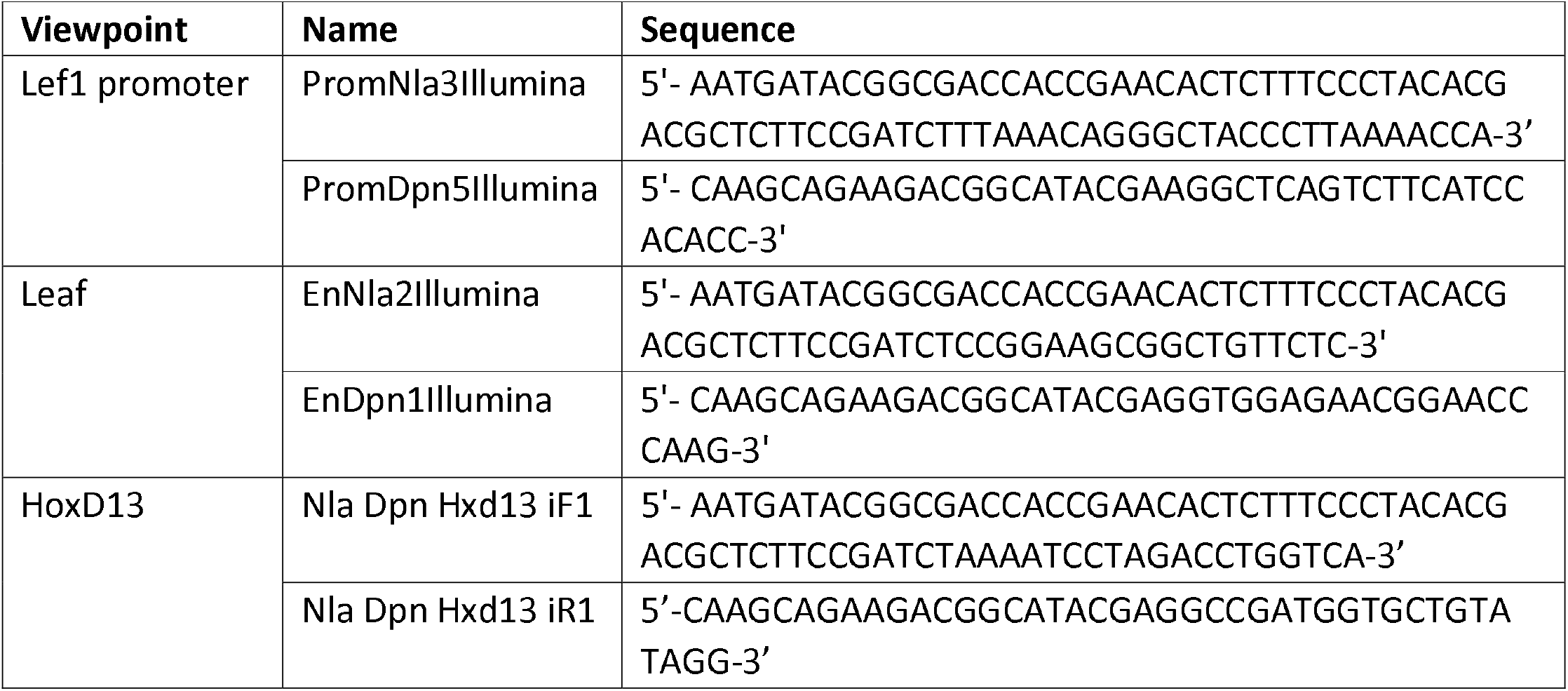
– List of 4C-Seq primers

## Materials and Methods

### Mice

OF1 and C57/B6J2 mice were obtained from Charles River Breeding Laboratories. *Prdm1*MEGFP transgenic mice were kindly provided by Mitinori Saitou; *Lef1* tm1 Rug mice were provided by Rudolf Grosschedl and Werner Held. *Sox2*Cre, *Sox2*CreERT2, *Prdm1*Cre, *Wnt1*Cre, *Prdm1* CA, ROSAYFP, mice were obtained from the Jackson Laboratory. The strains carrying the Cre recombinase and ROSAYFP were kept in heterozygosity, while the conditional knockout was performed in homozygosity. All animals were maintained in a 12⍰h light cycle providing food and water *ad libitum*. Mice were killed by intra-peritoneal (i.p.) injection of pentobarbital. Experiments were conducted in accordance with the EU Directive (86/609/EEC) for the care and use of laboratory animals and that of the Swiss Confederation. Mating of adult female and male mice was carried out overnight. Time-pregnant mice were killed by injection of pentobarbital and uteri with embryos were removed by dissection.

### Genotyping

List of primers is available in supplementary Table 1.

### *In situ* hybridization

RNA-FISH were performed using 8-μm paraffin sections. Digoxigenin-labeled probes for specific transcripts were prepared by PCR with primers designed using published sequences. The mRNA expression patterns were visualized by immunoreactivity with anti-digoxigenin horseradish peroxidase-conjugated Fab-fragments (Roche, Basel, Switzerland), according to the manufacturer’s instructions. The amplification was carried out using the TSA Plus Cyanine 3/5 System (Perkin Elmer). The probe SP72-Bmp4 and *Bmp2* were provided by Severine Urfer, Shh by the Duboule group, Lef1 by Anne Grapin-Botton. Gli1 and Wnt10b ISH were performed with the ACDBioscope.

### In vivo lineage tracing

For lineage tracing experiment, *Sox2*CreERT2/RosaYFP pregnant females were induced at embryonic day e12 and e12.5 with 2mg tamoxifen and 1mg of progesterone (Sigma-Aldrich) by intraperitoneal injection. The transgenic animals were retrieved at e17 and perinatally and then processed for histology and immunostaining.

### Proliferation experiments

For EdU experiment, pregnant female mice carrying *Prdm1*MEGFP embryos were injected with 200⍰µl of EdU (30⍰mg/⍰ml) and analyzed 2⍰h after the first injection. Embryos were retrieved, genotyped under the UV lamp and processed for histology and immunostaining.

### Histology and immunostaining

All samples were removed and fixed overnight in 4% paraformaldehyde at 4⍰°C. Tissues were washed three times in PBS for 5⍰min and incubated overnight in 30% sucrose in PBS at 4⍰°C; eventually they were then embedded in OCT and kept at −80⍰°C. Sections of 10⍰µm thickness were cut using a CM3050S Leica cryostat (Leica Mycrosystems).

Sections were incubated in blocking buffer (1% BSA, 0.3% Triton in PBS) for 1⍰h at room temperature. Primary antibodies were incubated overnight at 4⍰°. Sections were rinsed three times in PBS and incubated with appropriate secondary antibodies diluted to 1:1000 and DAPI in blocking buffer for 1⍰h at room temperature. Sections were again washed three times with PBS. The primary antibodies used are listed in supplementary table 2. The following secondary antibodies were used: anti-mouse, anti-rabbit, anti-rat, anti-goat, conjugated to Alexa Fluor 488, 568 and 647 (Molecular Probes). Nuclei were stained in DAPI solution (1:2000) and slides were mounted in DAKO fluorescent mounting medium. As for the AP reaction, SIGMA*FAST*^™^ Fast Red TR was used and visualized by confocal microscopy (Leica) at 568 nm.

### Imaging

Fluorescence microscopy images were captured under the LSM 780 confocal microscope (Carl Zeiss); transmission microscopy images were acquired with either the Olympus Ax70 or the Zeiss Axioscope 2 Plus.

### RT–PCR

Total RNA was isolated from the embryonic whisker pad using RNeasy Mini Kit (Qiagen) according to the manufacturer’s instructions. Total RNA was extracted, and 250 ng of each sample were reverse transcribed using the Superscript III enzyme and random primers (Life Technologies).

### Quantitative PCR

For qPCR, 1⍰µl of cDNA was amplified with the Taqman Universal Mastermix II (Life Technologies), in a 10⍰µl total reaction volume; 5µl of the Mastermix, 1,5µl of CDNA and 3,5µl of assay mix were included in the reaction. The primers were bought from Applied Biosciences or synthetized at IDT. The Taqman assays were performed using a 79000 HT Fast Real Time PCR system (AB). For data analysis, the mouse Eef1alpha, β-actin, Tbp and Gapdh housekeeping genes were used as internal control. Gene expression profiling was achieved using the Comparative CT method (DDC_T_) of relative quantification (Livak and Schmittgen, 2001) using the SDS 2.4 software (Applied Biosystems).

### Interspecies Sequence Comparison

The comparison of the conserved noncoding elements and deletions in mouse-rat, mouse-guinea pig, mouse-squirrel, mouse-rabbit, mouse-human, mouse-chimp, mouse-gorilla, mouse-orangutan, mouse-rhesus, mouse-dolphin, mouse-cow, mouse-cat, mouse-dog, mouse-horse, mouse-elephant was done using the Vertebrate Multiz Alignment and Conservation Track in the UCSC genome browser, using a window size of 2 kb.

### 4C-Seq

For each sample, we harvested at least 1×107 cells and obtained 7-10 μg of output double-digested, double-ligated DNA. We collected 120 embryonic whisker pads both at e12.5 and e13. NlaIII and DpnII were used as primary and secondary cutter respectively; ligation was performed by using the Concentrated T4 DNA ligase from Promega. Primer sets for *Lef1* promoter and enhancer are described in Supplementary table 2.

The primer set for *Hox*D13 has already been published. PCRs were multiplexed and sequenced with Illumina’s HiSeq2000. Raw data were subject to demultiplexing, mapping (mm10) and 4C analysis through the HTS station (http://htsstation.epfl.ch) according to previously described procedures. All figures were made using a running mean algorithm with a window size of eleven fragments. The regions excluded for analysis are for the promoter track chr3: 131104979-131112546 and for the enhancer tracks chr3: 131016310-131022769. Normalization was done by dividing the fragment scores by the mean of fragments scores falling into a region defined as +/- 1Mb (parameter) around the center of the bait coordinates.

### Statistical analysis

The normalized and profile corrected values were processed with Prism. For Leaf, fragments of region Chr3:131008663-131026430 (mm10) were analyzed whereas for the *Lef1* promoter and coding sequence, the fragments of region Chr3:131106987-131227057 were analyzed. Normality of the data was excluded by D’Agostino & Pearson omnibus normality test. Statistical differences were assessed by applying an unpaired non-parametric two-tailed Mann-Whitney test. Differences were regarded as significant if *p* < 0.05, and significances are shown in figures as * (* *p* < 0.05, ** *p* < 0.01, *** *p* < 0.001, **** *p* ≤ 0.001). Analyses were conducted using GraphPad Prism version 8.0.

## Acknowledgements

We thank Mitinori Saitou and Azim Surani for providing the *Prdm1*mEGFP mice, Rudolf Grosscheldl and Werner Held for providing the *Lef1*tm1Rug mice, Irene Pizzitola for breeding and mating *Lef1*tm1Rug mice, Michiko Kanemitsu for experimental support and constructive discussion, Giulio Cossu for the suggestions on the characterization of the pericytes and proofreading the manuscript, the EPFL CPG (Emilie Gesina, Gisele Ferrand) and the animal house of Epalinges (Francis Derouet, Lisa Arlandi) for mice handling, Orbicia Riccio, Elisabeth Joye and Severine Urfer for sharing protocols and probes for in situ hybridization, Andrea Zaffalon and Matteo Pluchinotta for molecular biology suggestions, Matteo Pluchinotta for artwork, Olga de Sousa Silva and Lai Quiewen for excellent technical help.

## Author contribution

P.G.M. conceptualization, experimental work, manuscript writing; F.D. conceptualization and experimental work on evolutionary part of the paper, manuscript writing; M.L. bioinformatic analysis; J.C., B.M., F.D, M.S. experimental work; G.F.M and H.A. histology, immunohistochemistry and ACDbioscope RNA-FISH; Y.B. conceptualization and supervision of the study.

