## Supplementary figures and images for "Early mechanisms of whisker development: Prdm1 and its regulation in whisker development and evolutionary loss"

### SFig1

Supplementary 1

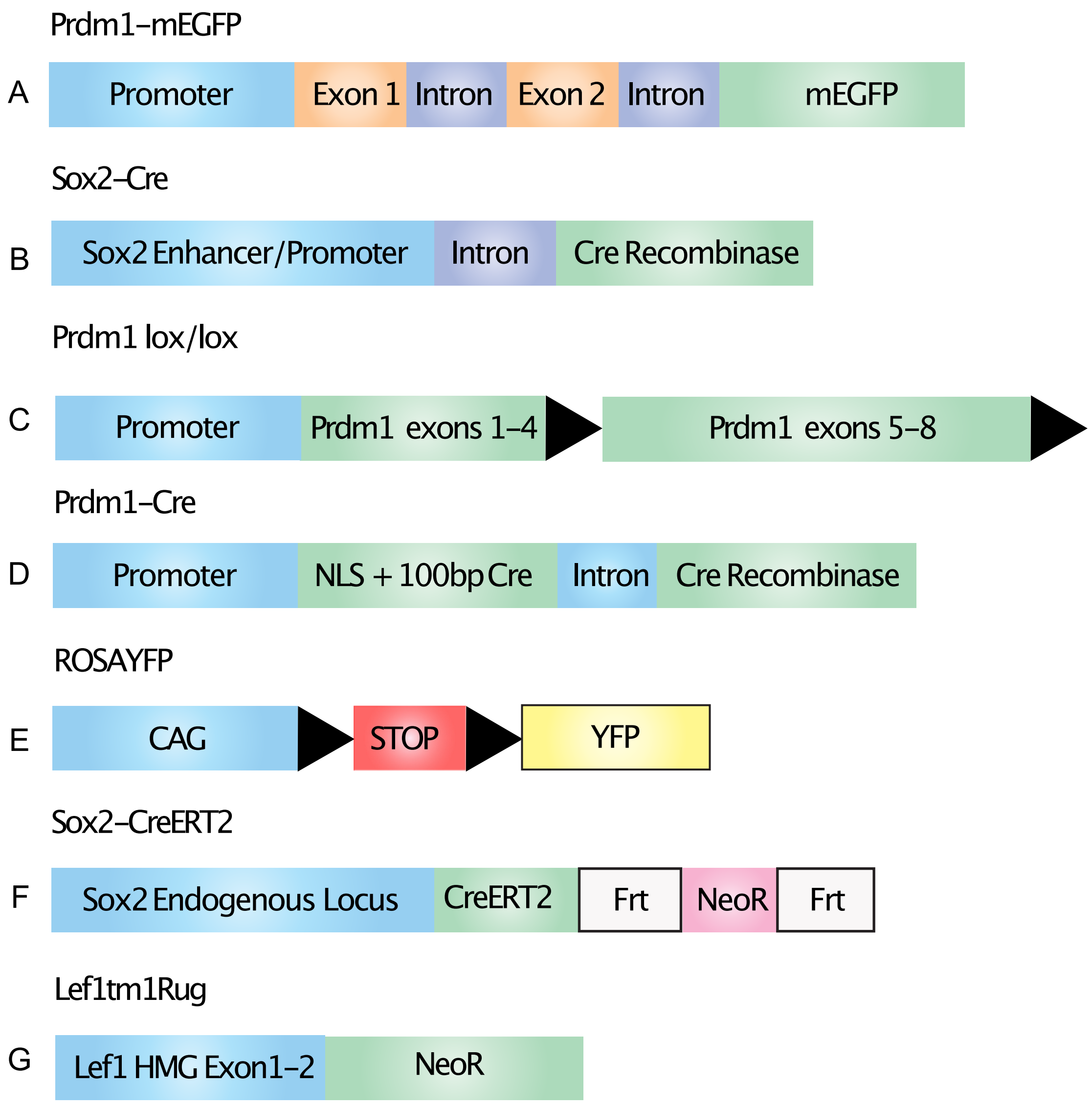

### SFig2

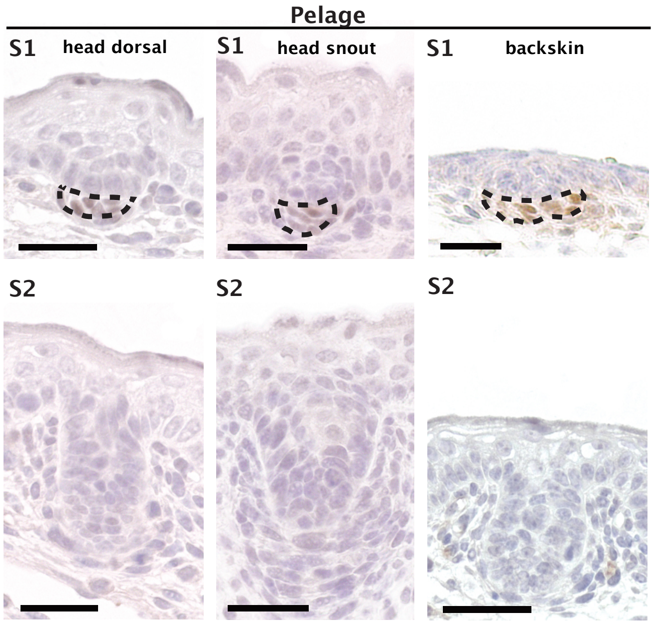

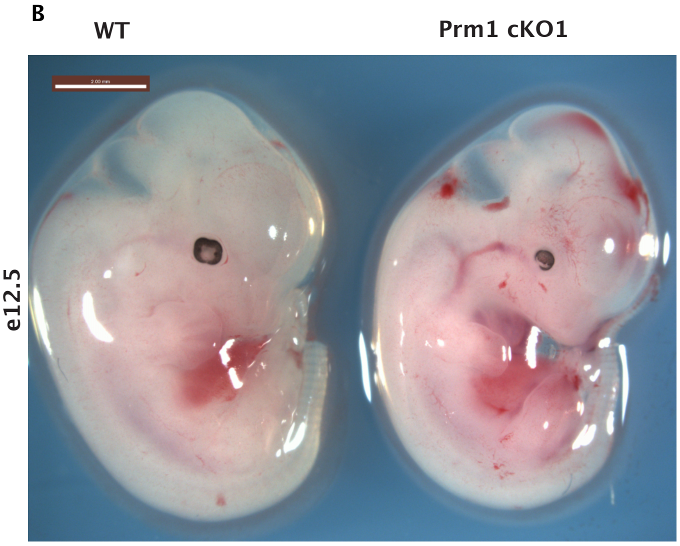

### SFig3

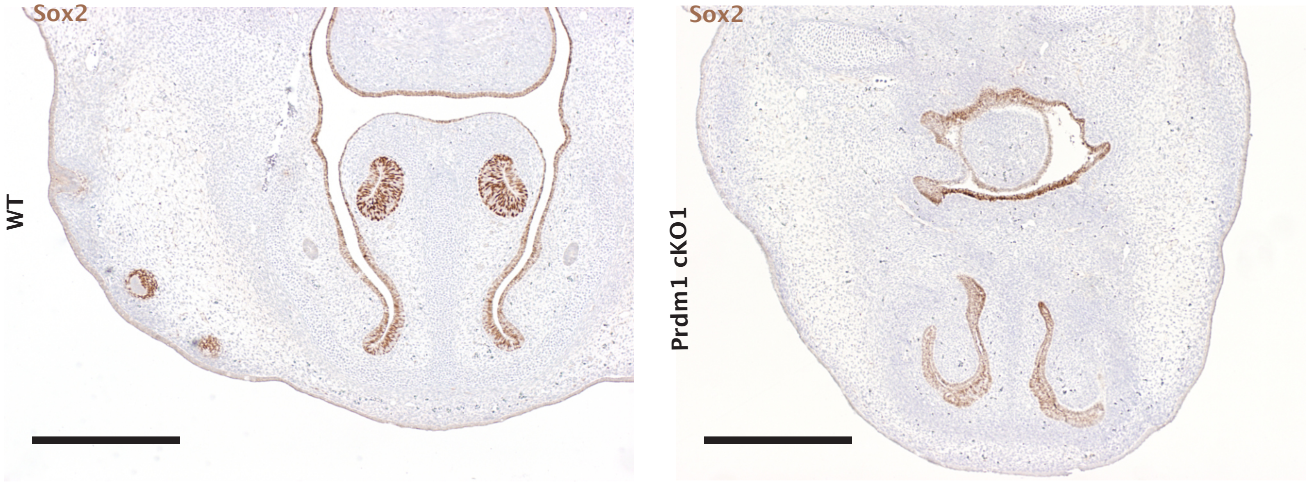

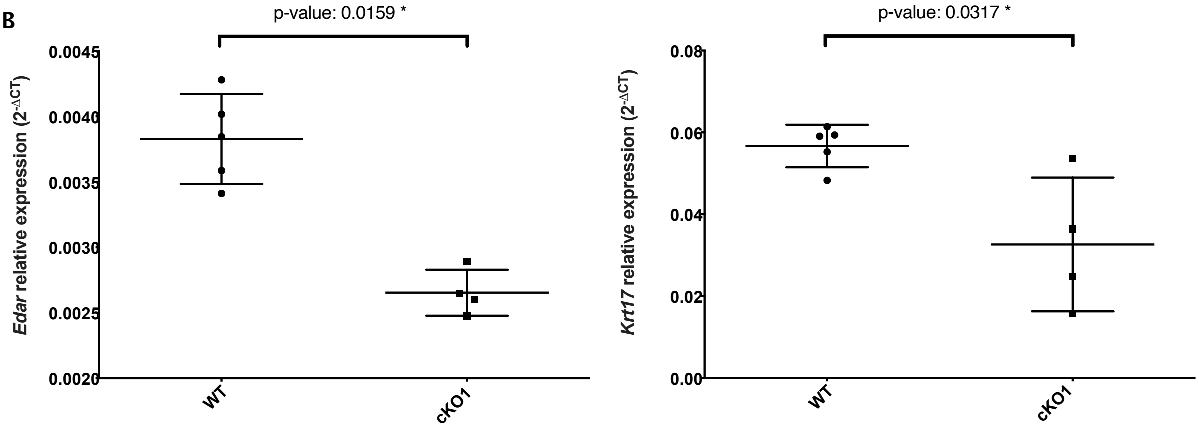

### SFig4

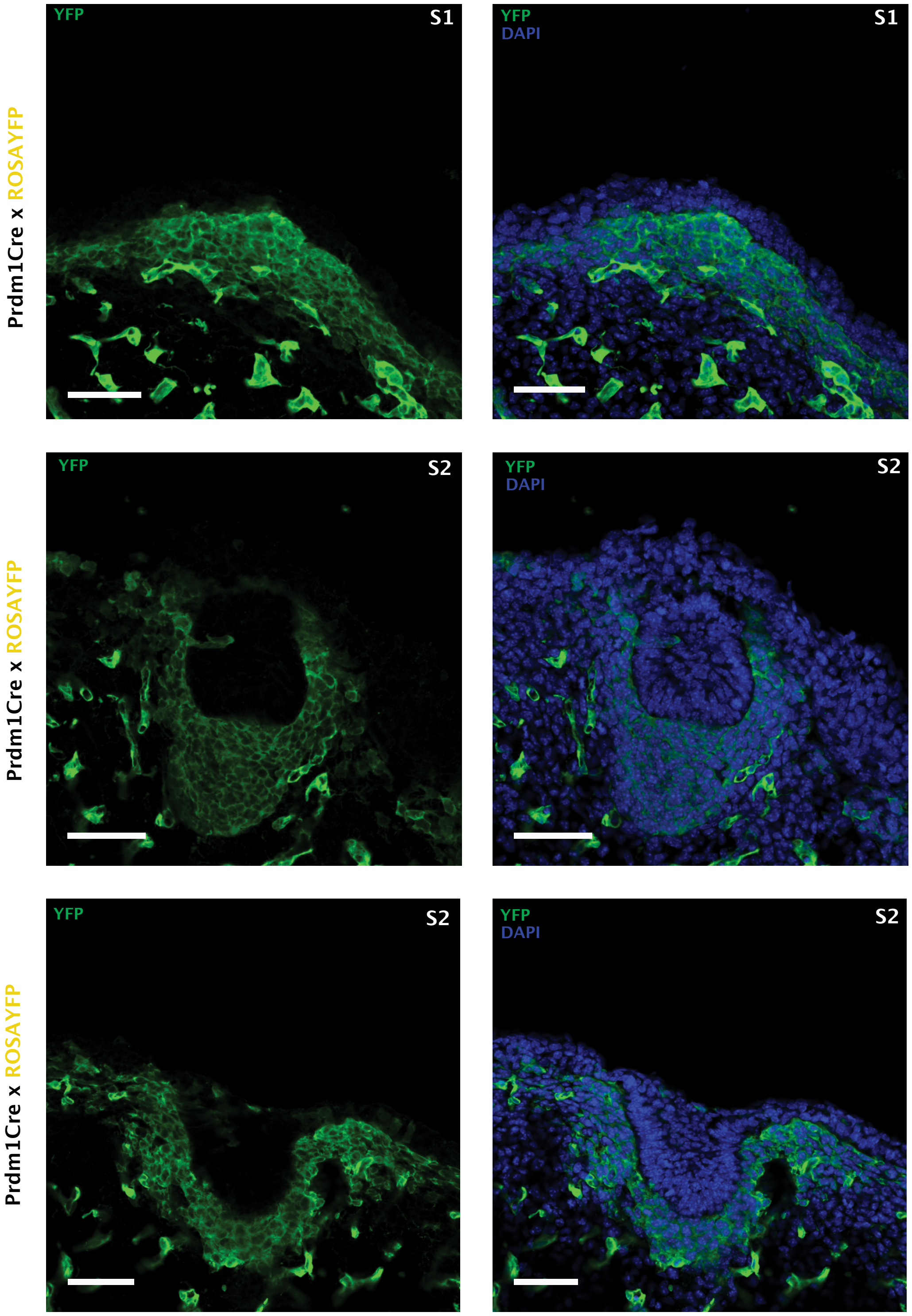

### SFig5

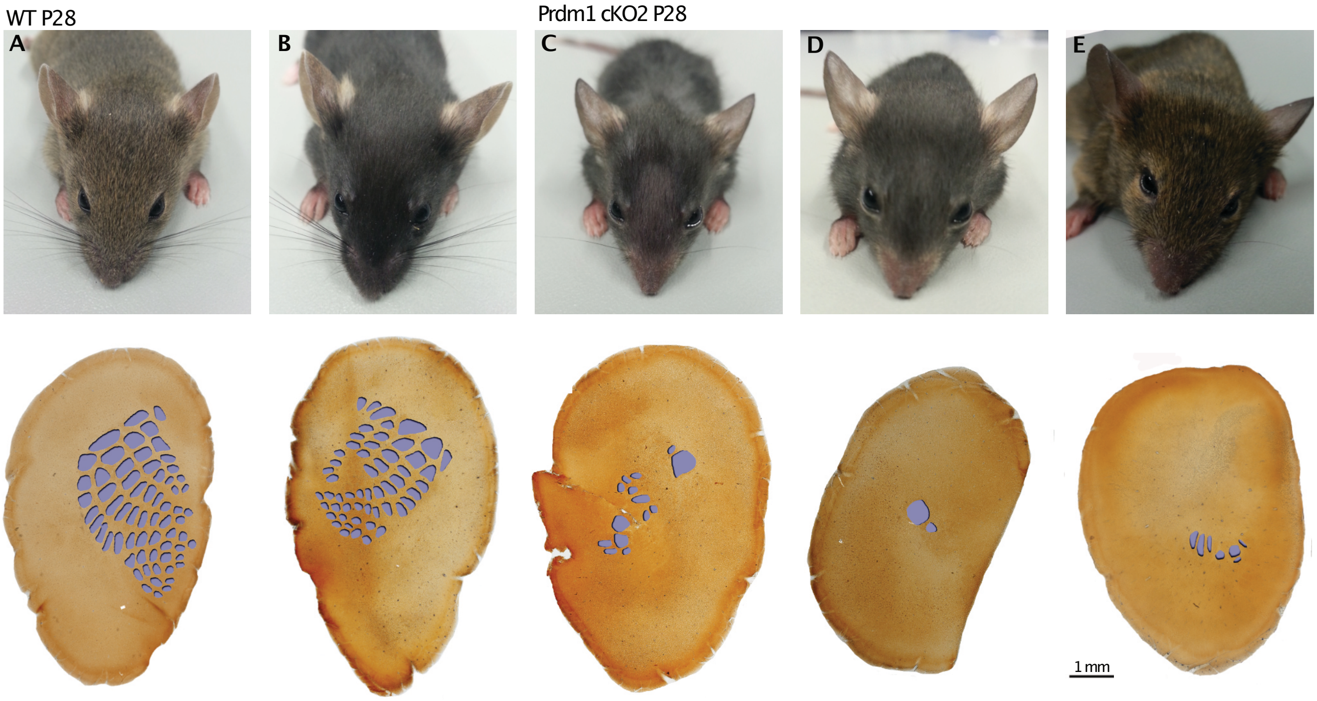

### SFig6

#### Supplementary 6

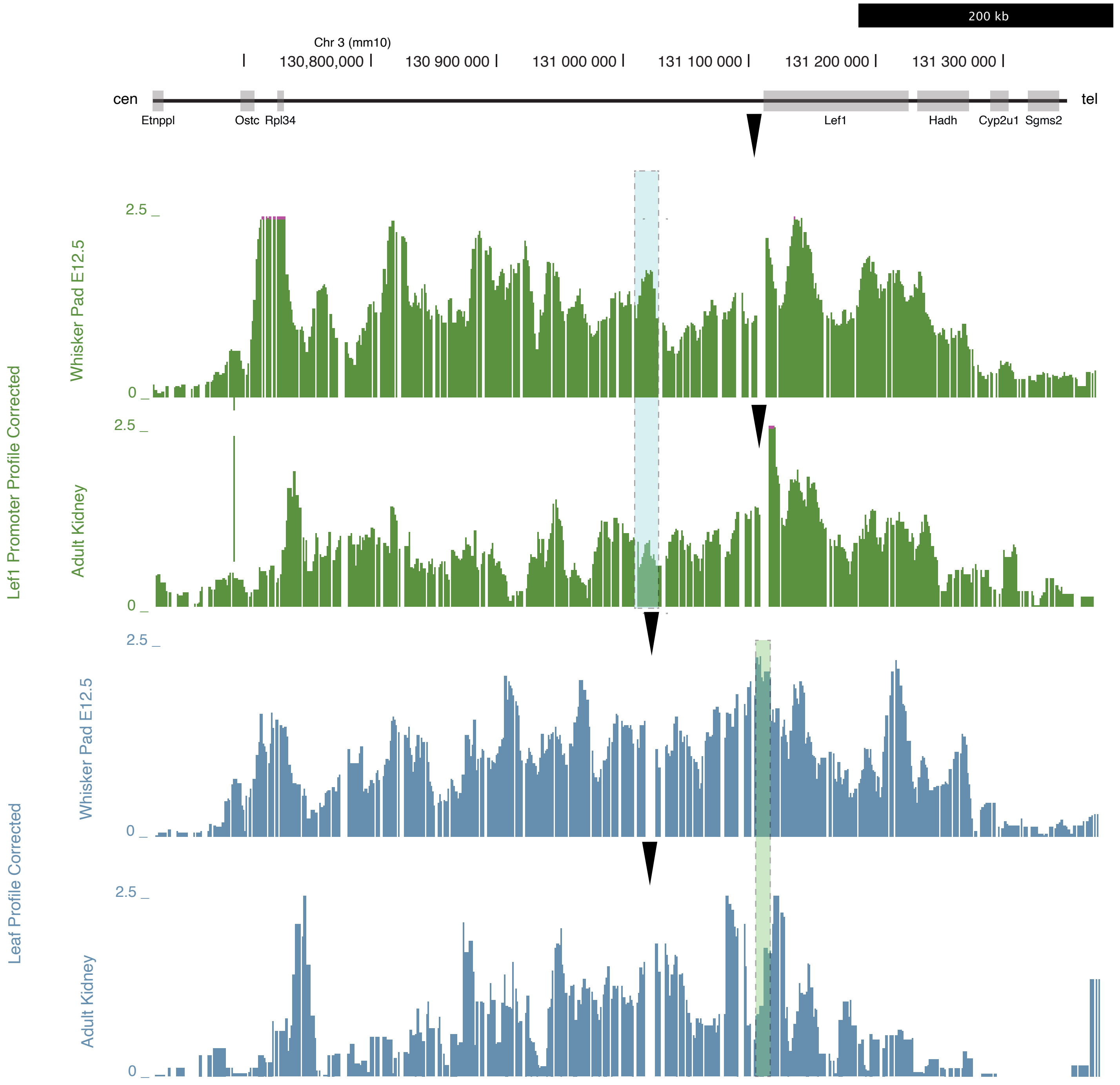
